# Discovery and evaluation of novel biomarkers reveal dasatinib as a potential treatment for a specific subtype of Triple-Negative Breast Cancer

**DOI:** 10.1101/2024.07.24.603752

**Authors:** D Ortega-Álvarez, D Tébar-García, M Casado-Pelaez, E Castillo-Agea, J Balibrea-Rull, D Olivares-Osuna, D Pérez-Parra, C Guardia, E Musulén, E Vinuesa-Pitarch, E Sánchez-López, F Postigo-Corrales, E Ballana, A Martínez-Cardús, M Margelí, M Esteller, PL Fernández-Ruiz, G Luengo-Gil, E Mereu, EM Galán-Moya, V Rodilla

## Abstract

Triple-negative breast cancer (TNBC) represents the most heterogeneous and aggressive subtype of breast carcinomas, characterized by the absence of clinical biomarkers (ERα, PR, and HER2) and the lack of targeted therapies. In this regard, several clinical trials have consistently failed to effectively stratify patients and identify specific treatments that elicit substantial responses. This study aims to pinpoint biomarkers expressed exclusively in the basal mammary epithelial compartment, facilitating a refined subclassification of this breast cancer subtype. Using computational analyses of single-cell RNA sequencing data, we have identified a list of genes associated with basal identity (BC-markers). Histological validation in 137 human samples has enabled us to categorize TNBC patients into BC-positive and BC-negative TNBC subtypes. Significantly, the presence of these markers correlates with a poorer prognosis in TNBC patients. Functional analyses have revealed a pivotal role for *TAGLN* in cell migration, likely influencing tumor aggressiveness. Further, we discovered that BC-marker expression is associated with the mesenchymal phenotype and increased sensitivity to the tyrosine kinase inhibitor dasatinib, particularly in BC-positive TNBC, suggesting novel therapeutic avenues. In our study, *TAGLN* emerged as a potential predictive biomarker for dasatinib responsiveness, offering new directions for personalized therapy for TNBC patients.

## Introduction

Breast carcinomas are clinically classified by the histological presence of hormone receptors, such as the estrogen receptor alpha (ERα) and progesterone receptor (PR), as well as overexpression of human epidermal growth factor receptor 2 (HER2) [1]. Tumors expressing ERα and PR are categorized as hormone receptor-positive (HR-positive), while those with elevated levels of HER2 are identified as HER2-enriched [1]. However, tumors devoid of these clinical biomarkers are diagnosed as triple-negative breast cancer (TNBC). Accounting for about 15% of all breast cancer cases, TNBC is notorious for its dire prognosis. This is attributed not only to its propensity for aggression and prevalence among younger women [2] but, crucially, because these patients lack targeted therapy options and must rely on chemotherapy as their standard of care [2].

Identifying new biomarkers and molecular targets for TNBC has been a daunting challenge due to its biological heterogeneity, which is evident in the variability of prognosis, pathological characteristics, treatment responses, and gene expression profiles [3]. The discovery of HER2 as a breast cancer marker exemplifies how identifying novel biomarkers can first recognize patients with different outcomes and, secondly, aid in developing new targeted therapies, such as monoclonal antibodies, small molecules, or CAR-T cells, among others. As such, the quest for molecular markers that can target specific tumor types has been a focal point of research. To understand breast tumor heterogeneity, several transcriptomic panels, such as the PAM50, which classifies breast cancer into luminal A/B, HER2-enriched, basal-like and normal-like subgroups [4–6], or the 70-gene *MammaPrint* microarray assay that has prognostic significance in ERα-positive and ERα-negative early-stage node-negative breast cancer [7], have been developed over the last decades. Specifically, for TNBC, Lehmann’s classification identified up to six distinct subgroups (Basal-like 1 and 2, Mesenchymal-like, Mesenchymal-stem like, Luminal androgen receptor and Immunomodulatory) [8]. These transcriptomic-based studies have clearly demonstrated that the heterogeneity of tumors has cemented the concept of breast cancer as a collection of different pathologies, each with varied clinical outcomes, rather than a singular disease [9,10]. Crucially, the clinical diagnosis of TNBC could be vastly enhanced through the discovery of novel molecular biomarkers that enable histological stratification, contributing to the development of new targeted therapies and significantly improving the prognosis of these patients.

The epithelial compartment of the mammary gland is composed of basal cells (BCs), which line the outer layer of the ducts and are known for their contractile ability, and luminal cells (LCs) that line the central lumen. LCs are subdivided into luminal ERα-expressing cells (ERα^pos^ LCs), hormone-sensing cells that can induce proliferation of the surrounding luminal ERα-negative cells (ERα^neg^ LCs), often referred to as hormone-responsive or alveolar cells [11]. Lineage tracing experiments in healthy murine mammary glands have shed light on the cellular hierarchy within this tissue. These studies have shown that in adult mice, different epithelial cell populations of the mammary gland are self-maintained by specific unipotent cells, indicating that these epithelial types contribute independently to mammary gland homeostasis by preserving their cellular identities into adulthood [12–14]. Recent single-cell RNA-sequencing (scRNA-seq) studies of the human mammary gland reveal similar epithelial cellular populations [15,16]. Notably, the molecular markers ERα and PR, currently used in breast cancer diagnosis, only positively identify ERα^pos^ LCs, while the other two epithelial cell types, ERα^neg^ LCs and BCs, remain histologically indistinct in clinical practice. Given that TNBC subtype consists of cells negative for these tested markers, it is intriguing to consider that defining new molecular markers that can distinguish each mammary epithelial cell type may significantly enhance our understanding of breast cancer heterogeneity. This could allow us to classify tumors with distinct cellular identities at histopathological level and characterize differences in their metastatic capacity and/or drug resistance, ultimately addressing two unmet clinical needs.

In this study, we have characterized novel molecular BC-associated markers (TAGL, SMA and TPM2) that play a crucial role in driving the malignancy of a specific subset of TNBCs. Remarkably, the expression of these biomarkers not only correlated with the aggressive nature of this breast cancer subtype but also significantly influences their sensitivity to clinically approved drugs, such as dasatinib. Dasatinib, a pan-tyrosine kinase inhibitor traditionally used in the treatment of certain hematological cancers [17,18], emerges as a promising therapeutic option for this specific subset of TNBC. Here, we propose using BC-associated markers to stratify TNBC patients, thereby identifying those who would benefit from dasatinib treatment. This approach represents a new direction in the personalized treatment of TNBC, offering hope for improved outcomes in a cancer subtype that has historically been challenging to manage.

## Results

### Identification of BC-markers exclusively expressed in the basal mammary compartment

Publicly available scRNA-seq data from three studies on murine healthy mammary glands (GSE109711 [19], GSE164017 [20], and GSE148791 [21]) were integrated to generate unique epithelial signatures capable of identifying three epithelial clusters corresponding to BCs and LCs (ERα^pos^ and ERα^neg^) (**Figure 1A, Supplemental Figure 1A-B**). Considering that approximately 77% of TNBC tumors are classified at transcriptomic level as basal-like by PAM50 [22], we first sought the differentially expressed genes (DEGs) between BCs and LCs, identifying 219 putative basal identity-associated genes (**Supplemental Figure 1C**). After selecting orthologues genes using the Human Protein Atlas (HPA), we compiled a Top 20 list of BC and LC-associated genes to distinguish both basal and luminal epithelial compartments (**Figure 1B, Supplemental Figure 2-3**).

**Figure 1.**
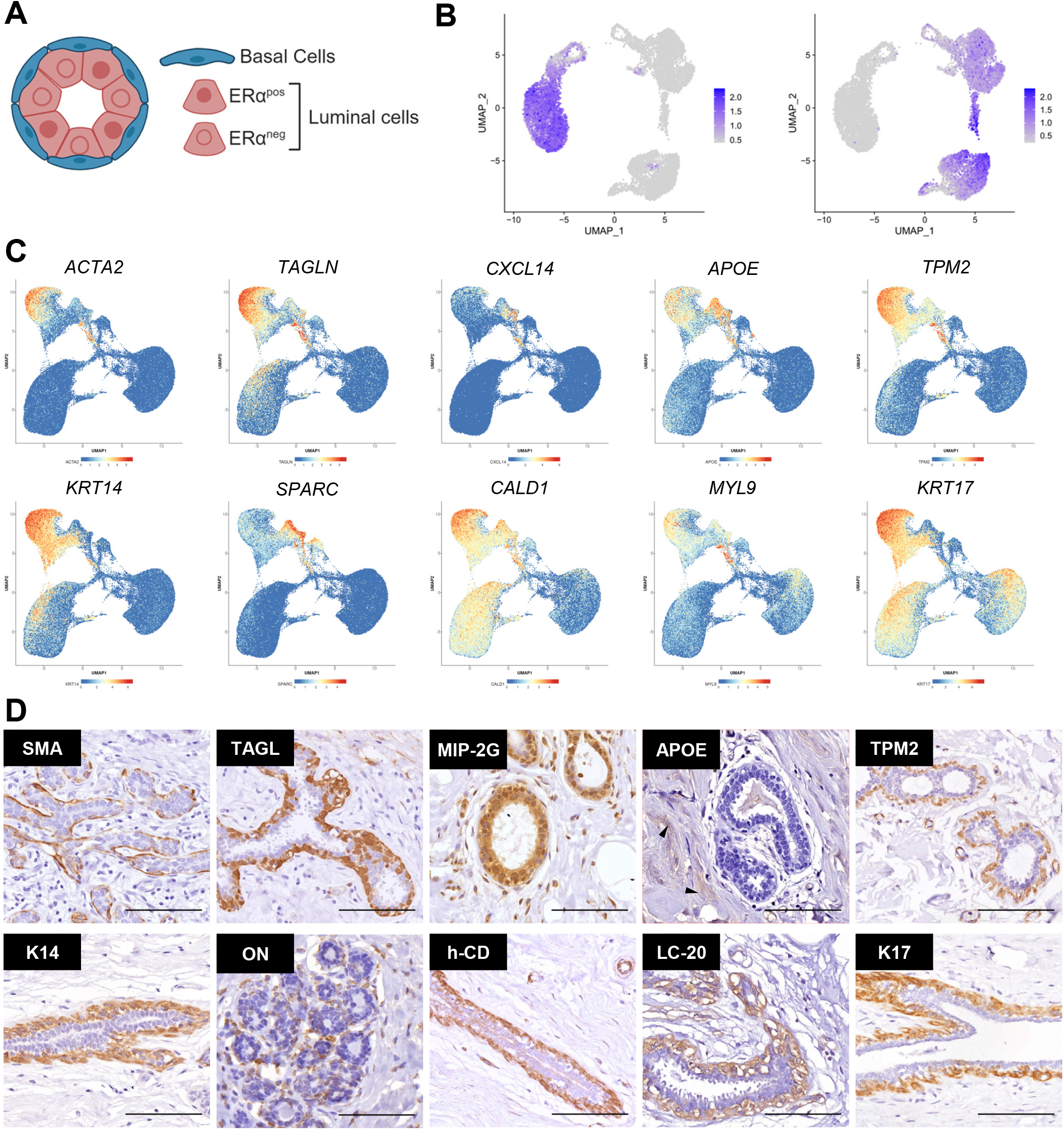
Identification of BC-identity makers expressed in the basal compartment of healthy human breast. **A**, Schematic representation of a cross-section of the mammary gland duct, composed of BCs in blue and LCs in red. ERαpos and ERαneg statuses are indicated by the color of the nuclei. **B**, UMAP plot showing the average expression of the Top-20 DEGs in the basal (left) and luminal (right) compartments from integrated scRNA-seq datasets of mammary epithelial cells from adult mice. **C**, Gene expression UMAP plots for Top-10 BC-associated genes from the integrated scRNA-seq datasets of healthy human breast tissue. Each plot depicts the expression level across cell populations, with a color gradient from blue (low expression) to red (high expression). **D**, Immunohistochemical staining panels showing the localization of proteins encoded by genes from panel C within the breast tissue. Scale bar represents 100 µm. Arrows indicate areas where the expression of APOE is restricted to the stromal compartment.

Subsequently, BC-markers were chosen based on their ability to consistently identify human BCs across both transcriptomic and immunohistochemical profiles. Three scRNA-seq studies performed on healthy human breast tissue (GSE161529 [16], GSE180878 [23], and GSE113197 [15]) were integrated (**Supplemental Figure 4**) to confirm the expression of BC-associated genes in human BCs. These analyses demonstrated that the Top 10 BC-markers were expressed in both murine and human BCs, reinforcing their utility as cross-species markers at transcriptomic level (**Figure 1C**). While the histological expression of some of these BC-associated markers, including Apolipoprotein E (Apo-E), C-X-C motif chemokine 14 (MIP-2G), and Osteonectin (ON), was not exclusively restricted to BCs within the mammary epithelial compartment (**Figure 1D**), protein-level validation confirmed exclusive expression of Smooth Muscle cell Actin (SMA), Transgelin (TAGL), β-tropomyosin (TPM2), Keratin 14 (K14), Caldesmon (h-CD), Myosin regulatory light polypeptide 9 (LC-20), and Keratin 17 (K17) in the BCs located in the outer layer of the human mammary epithelium in healthy breasts (**Figure 1D**). These results underscore the specific expression profiles that define the identity of the basal compartment in a healthy context.

### Specificity, diagnostic and prognostic value of BC-markers in TNBC

We evaluated the expression of seven BCs-markers in a cohort of 15 TNBC patients at diagnosis. Our results showed that SMA, TAGL, TPM2, LC-20 and h-CD were expressed in a specific subgroup of patients (2/15), which we have named BC-positive TNBC (**Figure 2A, Supplemental Figure 5-6**), representing about 13% of TNBC patients in this cohort. Notably, K14 and K17 were poorly expressed (<10% of positive cells) in the TNBC tumors analyzed (**Figure 2C, Supplemental Figure 5-6**), suggesting that the expression of these basal cytokeratin proteins is lost during the tumorigenic process.

**Figure 2.**
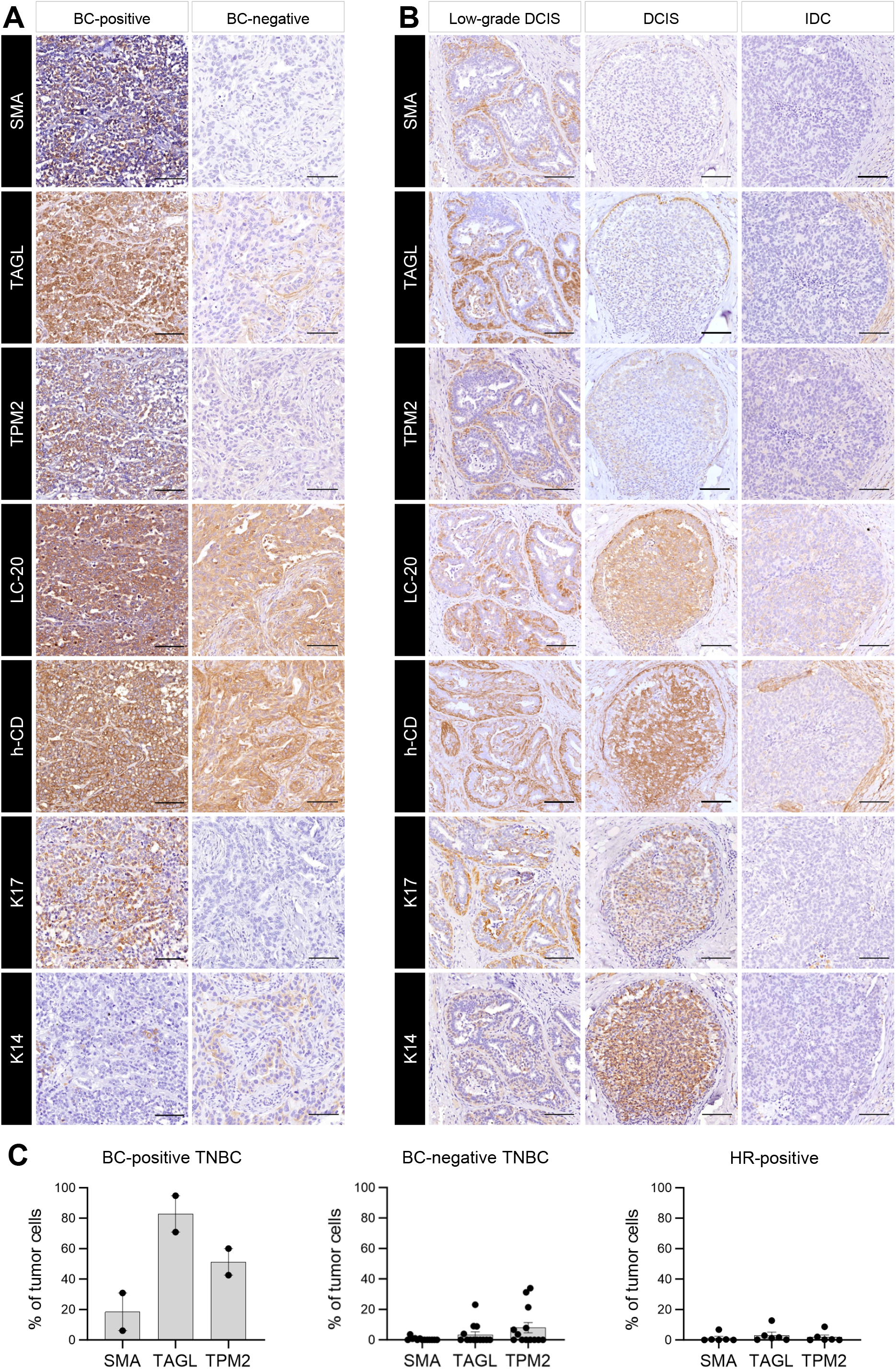
Histological expression of BC-markers in breast tumor samples. **A**, Representative sections of BC-positive and BC-negative TNBC samples stained with the indicated BC-markers. **B**, Representative sections of HR-positive samples at different tumor stages (low-grade DCIS, DCIS and IDC) stained with the indicated BC-markers. The scale bar represents 100 µm. **C**, Bar plots representing the percentage of tumoral cells expressing each marker in BC-positive TNBC, BC-negative TNBC and HR-positive samples.

The specificity of these BCs-markers was evaluated in HR-positive tumors (n=6) across different tumor stages, including low-grade ductal carcinoma in situ (low-grade DCIS), ductal carcinoma in situ (DCIS), and invasive ductal carcinoma (IDC). In low-grade DCIS, the expression of these exclusive BCs-markers was confined to the myoepithelial cells surrounding the tumor (**Figure 2B, Supplemental Figure 6A**), whereas in advance IDC samples, the expression of BC-markers was completely absent (**Figure 2B-C, Supplemental Figure 6C**), indicating that the basal layer of myoepithelial cells is lost during the progression of the HR-positive disease. In DCIS, the expression of SMA, TAGL, and TPM2 was exclusively present in the non-transformed myoepithelial cells, while LC-20, h-CD, K14 and K17 expression was also observed in tumor cells, suggesting that these BC-markers could be reactivated in LCs during tumor progression (**Figure 2B, Supplemental Figure 6B**). Consequently, LC-20, h-CD, K14 and K17 were excluded as exclusive BC-markers since they were not restricted to the BC-positive TNBC subtype, and their expression could also be observed in HR-positive tumors. After implementing this bottleneck strategy, we identified SMA, TAGL, and TPM2 as robust markers of basal identity. A computational analysis of the transcriptomic expression of these specific BC-markers in a dataset of TNBCs, using public databases (GSE81538 and GSE118527) [24,25], revealed that approximately 14% of TNBCs (9/63) could be classified as BC-positive subtype (**Figure 3A**). Additionally, RNA-seq analyses indicated that these BC-associated genes (*ACTA2, TAGLN*, and *TPM2*) were not distinctly associated with any particular subtype according to both the PAM50 (**Figure 3A**) and Lehmann classifications (**Figure 3B**).

**Figure 3.**
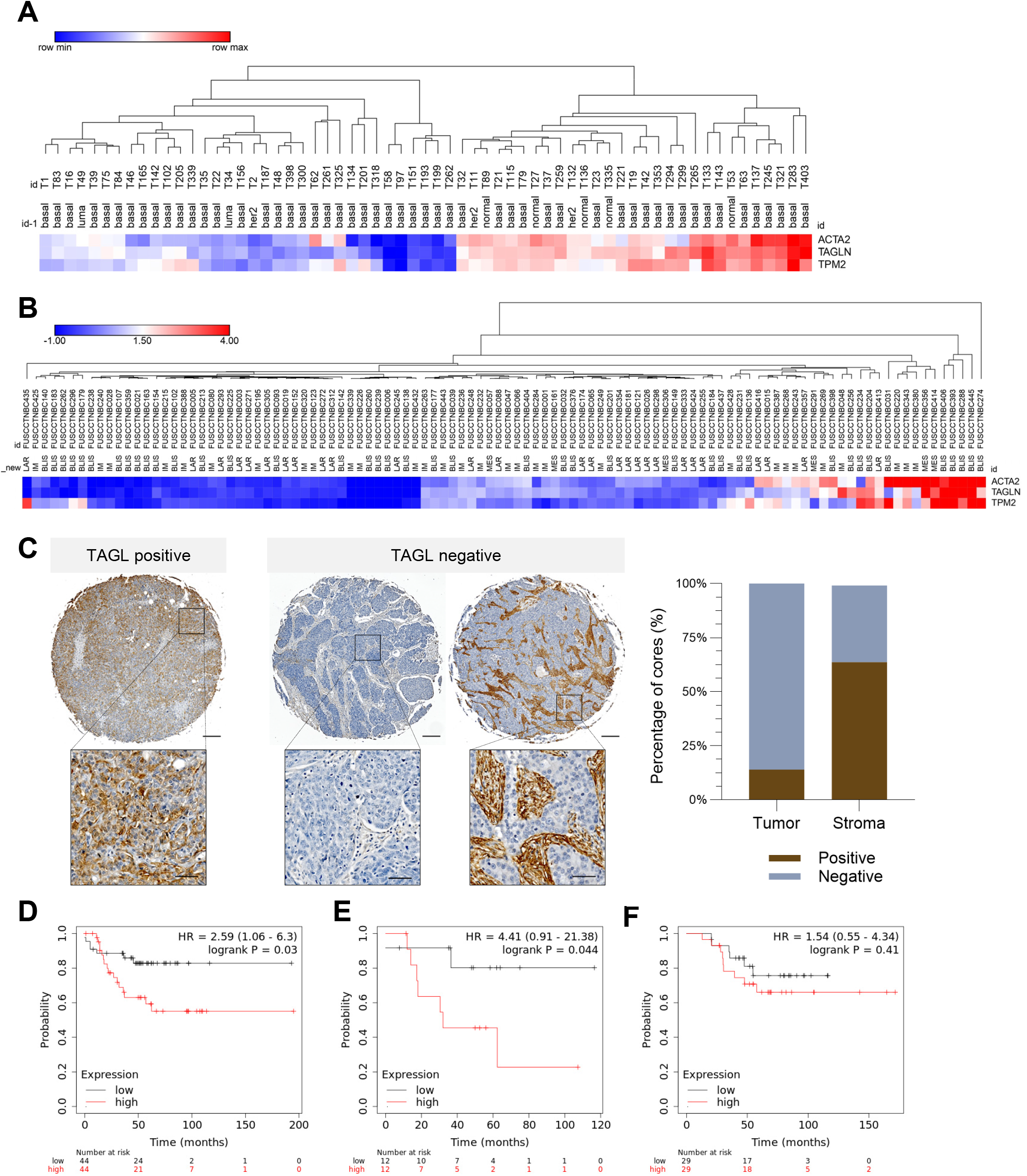
Expression and prognostic impact of BC-markers in TNBC. **A**, Heatmap of the transcriptomic profile of BC-markers in TNBC samples classified by PAM50. **B**, Heatmap of the transcriptomic profile of BC-markers in TNBC samples classified by Lehmann (high expression in red; low expression in blue). **C**, Representative cores of the tissue microarray showing TAGL staining in BC-positive and BC-negative tumors (left), and their corresponding quantifications on the right. Scale bar represents 200 µm and 50 µm in the insets. **D**, KM plot representing the recurrence free survival of TNBC patients based on BC-marker gene expression. **E**, KM plot representing the recurrence free survival of TNBC patients exclusively treated with chemotherapy based on BC-marker gene expression. **F**, KM plot representing the overall survival of TNBC patients based on BC-marker gene expression. Patient trichotomization was done comparing lower vs upper quartile (low expression in black; high expression in red).

To further evaluate the diagnostic potential of our BC-markers, we performed immunohistochemistry on a commercial TNBC tissue microarray containing n=122 samples. This analysis showed that 13,9% of the samples were positive for SMA, TAGL, and TPM2 (**Table 1, Figure 3C**), which is consistent with the percentage observed in our initial cohort of 15 TNBC patients. Notably, in BC-negative TNBC samples, the expression of these BC-markers was confined to stromal cells (**Figure 2A, 3C, Supplemental Figure 5**). These findings confirm that SMA, TAGL, and TPM2 serve as reliable and exclusive biomarkers for subclassifying TNBC, effective at both the transcriptomic and protein levels in pinpointing the BC-positive TNBC subgroup. The distinct expression patterns of these markers across various TNBC subtypes provide a comprehensive view of the molecular heterogeneity within this breast cancer subtype, highlighting their potential utility in identifying a distinct, previously uncharacterized group within the broader TNBC category.

**Table 1.** Distribution of clinical and pathological features in breast cancer samples. This table summarizes the expression of molecular markers and the clinical-pathological characteristics of TNBC patients, classified into BC-positive and BC-negative based on the staining for SMA, TAGL, and TPM2. It presents a breakdown of the percentage of patients expressing hormone receptors (ER/PR), HER2 status, and those with triple-negative profile (ER/PR/HER2-). The TNM classification is detailed along with nodal status and tumor grade. Each category is reported with the number of patients (N) and the percentage (%) relative to the total number of patients in the respective group. The total column reflects the sum of patients in both groups along with their percentage of the total cohort studied.

| Variable | BC-positive [N (%)] | BC-negative [N (%)] | Total |
| --- | --- | --- | --- |
| **Marker** |  |  |  |
| ER/PR (+) | 0 (0%) | 0 (0%) | 5 |
| HER2 (+) | 0 (0%) | 0 (0%) | 3 |
| ER/PR/HER2 (-) | 17 (13,9) | 105 (86%) | 122 |
| **TNM Classification** |  |  |  |
| T1 | 3 (30%) | 7 (70%) | 10 |
| T2 | 10 (13%) | 73 (87%) | 83 |
| T3 | 4 (17%) | 19 (83%) | 23 |
| T4 | 0 (0%) | 6 (100%) | 6 |
| **Nodal status** |  |  |  |
| N0 | 14 (17%) | 68 (83%) | 82 |
| N1/N2/N3 | 3 (7%) | 37 (93%) | 40 |
| **Tumor grade** |  |  |  |
| Grade 1 | 0 (0%) | 2 (100%) | 2 |
| Grade 2 | 8 (18%) | 35 (81%) | 43 |
| Grade 3 | 9 (11%) | 68 (88%) | 77 |

Then, to assess the clinical relevance of these novel BC-markers, we used the Kapplan-Meier (KM) plotter tool, which facilitates correlations between gene expression and survival curves across various tumor types. Our analysis comparing the mean expression of these specific BC-associated genes (*ACTA2, TAGLN*, and *TPM2*) between patients in the lower quartile and the upper quartile revealed that high expression of these markers is associated with a significantly worse prognosis, exhibiting a hazard ratio (HR) of 2.59 (**Figure 3D**). Moreover, patients treated exclusively with chemotherapy demonstrated an even poorer prognosis, with a HR of 4.41 (**Figure 3E**), suggesting that BC-positive TNBC patients do not benefit from standard chemotherapy regimens. Interestingly, no significant differences were observed in terms of overall survival, despite a HR of 1.54 (**Figure 3F**). These findings not only underscore the potential aggressiveness of BC-positive TNBC tumors compared to BC-negative ones but also highlight their resistance to conventional treatments, emphasizing the urgent need to explore alternative therapeutic options for this particular subgroup of TNBC patients.

### Exploring the tumorigenic role of TAGL in TNBC

Importantly, we investigated the tumorigenic role of BC-markers to determine whether they could serve as potential therapeutic targets. Given that TAGL exhibited the highest percentage of positive tumor cells in BC-positive TNBC samples (**Figure 2C**), we generated *TAGLN* knockout (KO) clones in two BC-positive TNBC cell lines (classified as mesenchymal-like), Hs578T and BT-549 (**Figure 4A-B**). Our findings indicated that *TAGLN* was not essential for cell proliferation, as evidenced by only a slight reduction in Hs578T cells and no significant differences in BT-549 cells following *TAGLN* depletion (**Figure 4C**). Since TAGL is an actin binding protein involved in multiple facets of cytoskeletal regulation [26,27], we assessed its impact on the migratory capacity of these TNBC cell lines *in vitro*. For *TAGLN*-KO clones, we observed a significant decrease in relative migration capacity compared to CONTROL cells in both in Hs578T and BT-549 cell lines (**Figure 4D**), suggesting that TAGL plays a crucial role in cell migration, which may influence breast cancer metastasis.

**Figure 4.**
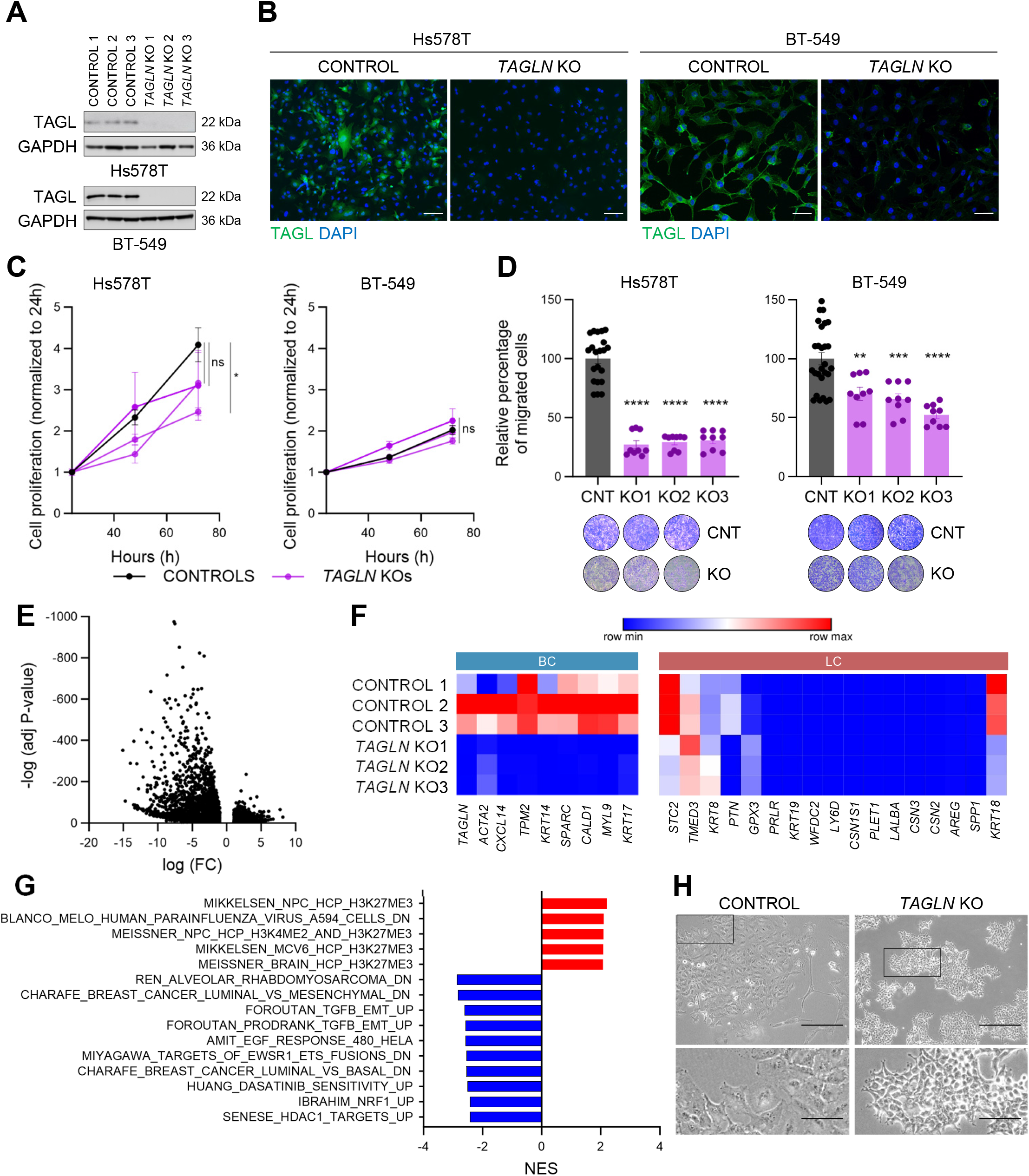
Functional validation of BC-markers in TNBC. **A**, WB of CONTROL and *TAGLN-*KO cells in Hs578T (above) and BT-549 (below) cells. **B**, Representative immunofluorescence staining in Hs578T and BT549 CONTROL and *TAGLN-*KO cells, stained with α-TAGL (green) and DAPI (blue). Scale bar represents 50 µm. **C**, Cell proliferation in CONTROL (black) and *TAGLN*-KO (purple) cells measured by MTT assays over 72 hours. **D**, Relative percentage of migrated cells in CONTROL (black) and TAGLN-KO (purple) cells, assessed by trans-well assays. In C and D, values represent the mean ± SEM of at least three experimental replicates, and statistical significance was determined using the *t test*. **E**, Volcano plot showing differentially expressed genes in *TAGLN*-KO cells compared to CONTROL Hs578T cells. **F**, Heatmap of the transcriptomic profile of BC and LC-associated genes in CONTROL and *TAGLN-KO* Hs578T cells (low expression in blue; high expression in red). **G**, Bar graph representing the normalized enrichment score (NES) of gene sets in *TAGLN-*KO compared to CONTROL Hs578T cells (positive NES in red, negative NES in blue). **H**, Brightfield images of Hs578T cells comparing CONTROL (left two panels) and *TAGLN*-KO (right two panels). Enlarged insets highlight morphological differences between the groups. The scale bar represents 300 µm and 100 µm in the magnification.

To further characterize the role of *TAGLN* in TNBC, we performed bulk RNA-seq on CONTROL and *TAGLN*-KO Hs578T cells, identifying a total of 2978 differentially expressed genes (**Figure 4E**). Notably, *TAGLN* deletion was accompanied by the downregulation of basal related genes, without inducing the luminal signature (**Figure 4F**). Using gene set enrichment analysis (GSEA), we identified key pathways altered in the absence of *TAGLN*, observing that among the most significant, epithelial-to-mesenchymal transition signatures were downregulated in *TAGLN-*KO cells (**Figure 4G, Supplemental Figure 7A**). This suggests that parental cells acquired an epithelial phenotype when *TAGLN* was depleted, as observed microscopically with changes in their morphology (**Figure 4H**), reinforcing the differences in their migratory capacity (**Figure 4D**). These observations align with our findings that BC-markers can identify a subset of TNBC with mesenchymal features (**Supplemental Figure 7A**), showing that TAGL plays an essential role in the aggressiveness of this BC-positive TNBC subtype, serving as a master regulator of basal/mesenchymal phenotype and regulating cell migration capacity, which could be crucial during the metastatic process.

### BC-marker expression and dasatinib sensitivity

Our data indicates that targeting TAGL may not be a viable therapeutic option, as no significant changes in proliferation were observed upon its deletion. However, TAGL has proven to be an excellent biomarker for stratifying patients, allowing us to explore novel therapeutic choices. With the aim of identifying targeted therapies for BC-positive TNBC patients, we observed a significant positive enrichment of genes related to dasatinib sensitivity in CONTROL compared to *TAGLN*-KO Hs578T cells (**Figure 4G, 5A**). Further analysis using publicly available data from the *DepMap* portal showed a positive correlation between dasatinib sensitivity and *TAGLN* expression in breast cancer cell lines (**Figure 5B, Supplemental Figure 8A**). To corroborate these findings, we conducted dose-response assays in a panel of TNBC cell lines (Hs578T, BT-549, MDA-MB-231, and HCC70) to establish a correlation between TAGL expression levels and sensitivity to dasatinib *in vitro* (**Figure 5C**). As a negative control, we used HR-positive cell lines (MCF7 and T-47D) which exhibited higher resistance to dasatinib (**Figure 5C**). Our results revealed that Hs578T cells showed the highest sensitivity to dasatinib (IC_50_ = 0,815 µM), followed by BT-549 (IC_50_ = 1,220 µM) (**Figure 5C**), suggesting that dasatinib could be a viable treatment option for BC-positive TNBCs, particularly those expressing high levels of TAGL. Additionally, we observed that all BC-markers analyzed in this study could potentially serve similar roles as biomarkers of dasatinib responsiveness (**Supplemental Figure 8C**). This suggests a broader utility of BC-markers in patients’ stratification to predict treatment efficacy, potentially extending beyond TAGL to include other markers in the panel.

**Figure 5.**
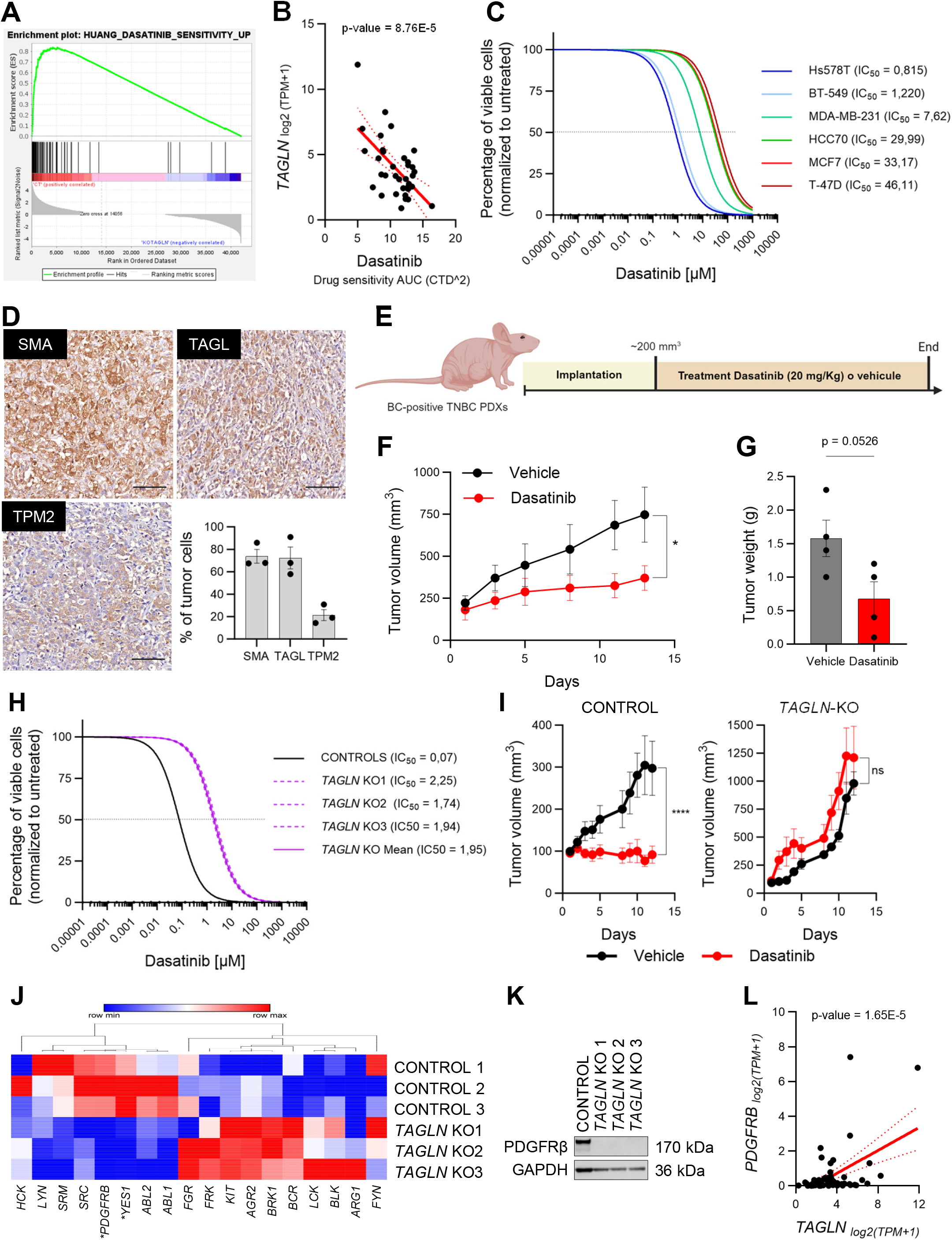
TAGL as a predictive marker for dasatinib sensitivity. **A**, Enrichment plot showing the GSEA for dasatinib sensitivity of CONTROL compared to *TAGLN*-KO Hs578T cells. **B**, Data from the *DepMap* portal showing the correlation between *TAGLN* expression (transcripts per million – TPM) and sensitivity to dasatinib (area under the curve – AUC); lower AUC indicates greater drug sensitivity. **C**, Drug-response curves for cell viability of indicated cell lines treated with dasatinib at the increasing concentrations. Solid lines represent the mean of three biological replicates performed in technical replicates, and error bars indicate ±SEM. The dashed line indicates their IC_50_. **D**, Representative sections of BC-positive PDX stained with the indicated BC-markers, with a bar plot showing the percentage of tumor cells per area expressing each marker. The scale bar represents 100 µm. **E**, Schematic representation of the dasatinib treatment protocol. **F**, Tumor volume of the BC-positive TNBC PDX treated with vehicle (black) or dasatinib (red). Error bars represent ± SEM. **G**, Bar plot showing the tumor weight of the BC-positive TNBC PDX treated with vehicle or dasatinib at the end point. Error bars represents ± SEM. **H**, Drug-response curves showing cell viability of CONTROL (black) and *TAGLN*-KO (purple) Hs578T clones treated with dasatinib at the indicated concentrations. Solid lines represent the mean of three biological replicates, and error bars denote ± SEM. **I**, Tumor volume measurements over days for CONTROL (left) and *TAGLN*-KO (right) Hs578T cells treated with vehicle (black) or dasatinib (red). Error bars represent ± SEM, with asterisks denoting statistical significance (***p<0.001, ns=not significant). **J**, Heatmap displaying the transcriptomic profiles of dasatinib target genes in Hs578T CONTROL and *TAGLN*-KO cells. Expression levels are color-coded, with blue indicating low expression and red indicating high expression. Asterisks mark genes with statistically significant differences in expression. **K**, WB analysis comparing the expression of PDGFRβ in Hs578T CONTROLs and *TAGLN*-KO cells. **L**, Scatter plots illustrating the correlation between *TAGLN* and *PDGFRB* expression in breast cancer cell lines. *TAGLN* expression (log2(TPM+1)) is plotted on the X-axis against *PDGFRB* expression (log2(TPM+1)) on the Y-axis. Black dots represent data points, with red dashed lines showing trend lines.

To further investigate whether BC-positive TNBC tumors could benefit from dasatinib treatment, we established a patient-derived xenograft (PDX) model. This PDX, positive for the three BC-markers (**Figure 5D**), was treated with dasatinib or vehicle daily for two weeks, with tumor growth measurements taken every other day (**Figure 5E-F**). Our results demonstrated that BC-positive TNBCs were significantly sensitive to dasatinib *in vivo*, as shown by reduced tumor growth and weight in dasatinib-treated mice compared to controls (**Figure 5F-G**). This finding underscores the potential of dasatinib as an effective therapeutic option for a subset of TNBC patients characterized by high expression of specific BC-markers.

### The role of *TAGLN* in dasatinib responsiveness

To elucidate whether *TAGLN* expression directly influences the response to dasatinib, we performed dose-response assays treating both CONTROL and *TAGLN*-KO cells with various concentrations of the drug. The results revealed that the abrogation of *TAGLN* conferred resistance to dasatinib, as evidenced by an increase in IC_50_ values compared with control clones in both Hs578T and BT-549 cells (1.95 vs 0.07 µM, and 4.315 vs 0.74 µM, respectively) (**Figure 5H, Supplemental Figure 7B**). Furthermore, we investigated the role of *TAGLN* in response to dasatinib *in vivo* by injecting CONTROL and *TAGLN*-KO Hs578T cells into the mammary fat pad of immunodeficient mice, followed by dasatinib treatment for one week. These experiments demonstrated that tumors derived from *TAGLN*-expressing cells were more susceptible to dasatinib, confirming the *in vitro* findings and suggesting a crucial role for *TAGLN* in mediating dasatinib sensitivity (**Figure 5I**).

Given the observed increase in sensitivity to dasatinib among TNBC cells expressing *TAGLN*, we aimed to elucidate the underlying molecular interactions responsible for this response. We found that several known dasatinib targets, including *ABL1, ABL2, YES1, PDGFRB, SRC, SRM* and *LYN*, were upregulated in CONTROL versus *TAGLN*-KO Hs578T cells (**Figure 5J**), however only *YES1* and *PDGFRB* were statistically significant. Importantly, the validation at protein level showed that among all the known dasatinib targets, only PDGFRβ expression was lost in *TAGLN-*KO cells compared to the CONTROL clones (**Figure 5K**), suggesting a critical role of TAGL in the regulation of the expression of this receptor. We identified a positive correlation between *TAGLN* expression and these key signaling molecules in breast cancer cell lines (**Figure 5L, Supplemental Figure 8D**). These results not only shed light on the molecular mechanisms underlying dasatinib sensitivity in TNBC but also establish *TAGLN* as a crucial component of the signaling pathways that enhance drug responsiveness.

## Discussion

Triple-negative breast cancer is a highly heterogeneous disease with variable prognoses and responses to treatment. The identification and characterization of novel biomarkers in this study have significant implications for improving the clinical management of TNBC, particularly in subclassifying tumors more accurately and tailoring therapeutic strategies. Our findings that BC-markers can stratify TNBC into at least two distinct subtypes, BC-positive and BC-negative, highlight the potential for more personalized treatment approaches. Importantly, the BC-positive subtype, characterized by the expression of markers such as *TAGLN*, showed a poor response to standard chemotherapy but increased sensitivity to the clinically approved tyrosine kinase inhibitor dasatinib, suggesting a pivotal shift in treatment strategies could be warranted. The demonstration that *TAGLN* and other BC-markers are not merely expressions of cellular identity but are also integral to the signaling pathways that govern tumor aggressiveness and treatment responses sheds light on the complex biology of TNBC. The association of these markers with dasatinib sensitivity, particularly through the modulation of Src family kinases and other dasatinib targets, underscores the potential for these markers to guide the use of targeted therapies in a clinical setting. Crucially, it has been demonstrated a direct correlation between TAGL and Src activation in ovarian cancer [28]. Specifically, ovarian cancer cells treated with dasatinib exhibited a substantial reduction in TAGL expression and Src activation, and the silencing of *TAGLN* also led to decreased levels of Src activation. Conversely, ectopic expression of Src active forms or TAGL overexpression increased TAGL and Src activation, respectively [28]. Considering our findings, this suggest that TAGL may participate in a regulatory loop of several dasatinib-associated targets serving as a predictive biomarker for its response.

Our study underscores the critical role of integrating scRNA-seq data and protein-level validation to identify and validate the functional roles of novel biomarkers. This comprehensive approach has enabled us to pinpoint markers that are diagnostic and prognostic, offering a richer understanding of tumor biology and revealing potential therapeutic targets. In our scRNA-seq analysis of healthy human breast samples, we observed that several genes (*MYL9, CALD1, KRT14* and *KRT17*) were not exclusively expressed in BCs, and interestingly, these were the same markers that were not exclusively found in BC-positive TNBC samples. The reactivation of some of these genes in luminal tumors suggests a potential acquisition of undifferentiated traits, reminiscent of those seen in the mammary bud during embryonic development. Notably, K14 expression has been detected in both epithelial compartments (LCs and BCs) in aged women, highlighting a potential age-related shift in gene expression patterns [23,29]. In the context of tumors, research has shown that low-ERα tumors may begin expressing basal cytokeratin proteins during the tumorigenic process [30]. Additionally, collective invasion of luminal tumors has been found to require K14-expressing cells [31], suggesting a critical role for these markers in tumor progression and metastasis.

Considering current knowledge on transcriptomic classifications of breast cancer, our BC-markers (*ACTA2, TAGLN*, and *TPM2*) are not associated to any specific TNBC subtype according to PAM50 and Lehmann classifications. It is important to note that basal-like tumors (by PAM50 and Lehmann) typically exhibit an enriched expression of classical basal cytokeratin proteins such as K5, K14, and K17. However, we have demonstrated that these proteins are not specific to BC-positive tumors. This observation suggests that basal-like breast tumors may actually represent luminal TNBC tumors that acquire basal cytokeratin expression during tumor progression.

Furthermore, our research indicates that the mesenchymal-like features of the BC-positive subtype may be key drivers of the aggressive nature and chemoresistance observed in these tumors. The ability of *TAGLN* to regulate EMT and maintain mesenchymal characteristics suggests that targeting this pathway could inhibit tumor progression and metastasis. The correlation between high *TAGLN* expression and poor prognosis reinforces the need for targeted therapeutic strategies that could disrupt these signaling pathways, offering hope for improved outcomes in TNBC patients. Interestingly, the role of *TAGLN* in tumor progression was also explored in other tumor types, including its involvement in cell proliferation, migration, and inhibition of tumor formation in colorectal cancer [26], non-small cell lung cancer [32] and lung adenocarcinoma [33].

Looking ahead, the application of BC-markers in clinical trials would be crucial to validate these findings and potentially integrate them into routine diagnostic and therapeutic protocols. Moreover, investigating the interaction between BC-markers and the immune landscape of TNBC may uncover synergies with immunotherapies, which have been effective in other breast cancer subtypes. Importantly, the observed correlation between BC-marker expression and dasatinib sensitivity is promising but requires validation in clinical trials. To this end, two clinical trials (NCT00817531 and NCT00371254) have assessed the efficacy of dasatinib in TNBC, demonstrating a clinical benefit rate of only 10%, which aligns closely with the prevalence of BC-positive tumors. These findings suggest a nuanced response to dasatinib, potentially limited to a specific subset of TNBC patients. Our results could allow a better stratification of TNBC patients that would benefit from this treatment.

In conclusion, our study provides compelling evidence that BC-markers, particularly *TAGLN*, hold significant promise as tools for improving the precision of TNBC diagnosis and treatment. By delineating the molecular characteristics of TNBC subtypes, we pave the way for more effective and individualized therapies, aligning with the broader goals of precision medicine in oncology.

## Methods

### Single cell RNA-sequencing analysis

We initially preprocessed each scRNA-seq dataset independently, including murine healthy mammary glands (NCBI Gene Expression Omnibus [GEO] accession numbers GSE109711, GSE164017, GSE148791) and healthy human breast tissue (GEO accession numbers GSE161529, GSE180878, GSE113197). The original count matrices were analyzed with Seurat (version 4.0.4) in R (version 4.0.3). Low-quality cells were filtered based on mitochondrial RNA content (>25%) and the number of genes expressed, excluding cells with gene counts below the threshold set between the minimum and the first quartile of the gene count distribution for each dataset. Each dataset was normalized by total expression, scaled by a factor of 10,000, and log transformed. Gene expression measurements were then scaled and centered. Highly variable genes (HVGs) were selected by assessing the relationship between gene dispersion and log mean expression. Principal component analysis (PCA) was performed, and the optimal number of principal components was selected by inspecting the elbow plot. We retained the top 20 principal components for downstream analysis. Dimensionality reduction was achieved using the Uniform Manifold Approximation and Projection (UMAP) algorithm. Cluster analysis was performed using the Louvain clustering algorithm. Clusters were annotated based on the expression of canonical markers to consistently identify luminal and basal populations. Additionally, for some datasets, prior cell-level annotation information was also available. To integrate the datasets (separately for murine and human samples), we used the raw counts from the previously annotated datasets and followed Seurat’s canonical correlation analysis (CCA) integration pipeline. This method identifies anchor cells between pairs of datasets, which are then used to harmonize the integrated datasets. Marker genes for each cell type were identified using Seurat’s *FindAllMarkers* function, considering only positive markers and setting min.pct to 0.25. Gene signature scoring was performed using Seurat’s *AddModuleScore* with default parameters.

### Human samples

In this study, two distinct cohorts of breast cancer samples were utilized. The first cohort comprised a total of 6 HR-positive and 15 TNBC samples from the Hospital Germans Trias i Pujol and the General University Hospital of Albacete. These samples were collected with the ethical approval of our research campus committee (CEIC #PI-21-230). Additionally, we used a commercial TNBC tissue array (#BR1301a), which included 122 cases with corresponding cores (one core per case). This array provided detailed information on pathology grade, TNM classification, clinical stage according to the AJCC 8th edition, and immunohistochemistry results for ERα, PR, and HER2.

### Immunohistochemistry (IHC)

Formalin-fixed and paraffin-embedded samples, sectioned at 4 μm, were dewaxed using a graded series of ethanol. Antigen retrieval was carried out by boiling the sections for 20 minutes in either citrate buffer solution (Sigma-Aldrich, #C9999) or Tris-HCl buffer solution (Sigma-Aldrich, #T6455). To block endogenous peroxidase activity, sections were incubated for 10 minutes in a 3% hydrogen peroxide solution. Permeabilization and blocking were achieved using a blocking buffer containing 0.3% of Triton ® X-100, 2% BSA and 5% FBS. Sections were then incubated overnight at 4ºC with primary antibodies diluted in the blocking buffer at the concentrations indicated in **Supplemental Table 1**. This was followed by 1-hour room temperature incubation with secondary antibodies (**Supplemental Table 2**). Sections were developed using Signal Stain DAB Substrate Kit (Cell Signaling Technology, #8059S) according to the manufacturer’s instructions and counterstained with Hematoxylin. Mounting was performed using DPX mounting solution for histology (MERCK, #06522). Whole tissue images were captured with Pannoramic SCAN II and analyzed using QuPath software. Tumor areas were delineated by a pathologist in images using the Brightfield (H-DAB) format, and a positive cell detection algorithm was applied to identify positive and negative cells and to quantify the percentage of positive cells within the selected areas.

### Cell culture

TNBC cell lines HS578T (DSMZ, #ACC781), BT-549 (ATCC, #HTB-122) MDA-MB-231 (ATCC, #CRM-HTB-26) and HCC70 (ATCC, #CRL-2315), along with HR-positive cell lines MCF7 (DSMZ, #ACC115) and T-47D (DSMZ, #ACC739), were cultured in DMEM/F-12 with GlutaMAX™ Supplement medium (Gibco™, #31331093), supplemented with 10% FBS, at 37ºC and 5% CO_2_ atmosphere. For the CRISPR/Cas9 knockout transfection, Hs578T and BT-549 cells were co-transfected with a ribonucleoprotein (RNP) complex comprising Cas9 and guide RNA (gRNA) according to manufacturer’s instructions (Invitrogen™, #CMAX00015). The crRNA and tracrRNA (Integrated DNA Technologies, #1072534) were mixed at equimolar concentrations, heated at 95ºC for 5 minutes, and allowed to cool at room temperature. To assemble the RNP complex, Cas9-GFP enzyme (Integrated DNA Technologies, #10008161) was prepared at 1 μM in Opti-MEM™ Reduced Serum Medium (Gibco™, #31985047). Each gRNA was then combined with the Cas9-GFP in Cas9 PLUS™ Reagent and Opti-MEM. The RNP complex was subsequently mixed with CRISPRMAX™ transfection reagent (Invitrogen™, #CMAX00015) at room temperature for 20 minutes. After 24 hours at 37ºC, transfected cells were plated in 96-well dishes for cellular subcloning.

### Immunofluorescence (IF)

The Hs578T and BT-549 cells were fixed with a 4% PFA solution for 10 minutes at room temperature. Permeabilization and blocking were performed using a blocking solution containing 0.3% Triton® X-100, 2% BSA, and 5% FBS for one hour at room temperature. Cells were then incubated for one hour at room temperature with primary antibodies diluted in the blocking solution at the concentrations indicated in **Supplemental Table 1**. This was followed by incubation with secondary antibodies (**Supplemental Table 2**), for one hour at room temperature. Mounting and DNA/nucleus staining were subsequently performed using Fluoroshield™ mounting medium with DAPI. Images were captured using an EVOS M500 microscope and processed with Fiji-ImageJ software.

### Western blot (WB)

For protein extraction, cells were pelleted and washed with cold PBS before being lysed in RIPA buffer (Thermo Scientific™, #89900) supplemented with 10 µL/mL Halt Protease and Phosphatase Inhibitor Cocktail EDTA-Free (Thermo Scientific™, #78440). Following centrifugation at 14,000g for 15 minutes, the supernatant was collected to quantify protein concentration using the BCA Pierce protein assay. Prior to electrophoresis, samples were heated at 95°C for 5 minutes. Running Buffer 1X was prepared by diluting MES SDS Running Buffer 20X (Thermo Scientific Chemicals, #J62138.K2). A precast gel was placed in the tank and filled with Running Buffer 1X. Samples were loaded into the gel and electrophoresed at a constant voltage of 200V. Protein transfer was performed using the iBlot™ Gel Transfer Device (Invitrogen™, #IB21001), according to the manufacturer’s instructions. Membranes were blocked in a buffer containing 5% milk, 0.1% Tween-20 in PBS for 1 hour at room temperature. After blocking, membranes were incubated overnight at 4°C with primary antibodies (**Supplemental Table 1**) diluted in 5% BSA and 0.1% Tween-20 in PBS with shaking. The next day, membranes were incubated with secondary antibodies (**Supplemental Table 2**) for 1 hour at room temperature. For protein detection, ECL solution (Thermo Scientific™, #32109) was used, and images were captured using a Vilber Smart Imaging Fusion FX developer.

### Proliferation assay

To conduct the CellTiter 96® Non-Radioactive Cell Proliferation Assay (MTT) (Promega, #G4000), 5,000 cells were seeded in 100 µL of DMEM/F-12 with GlutaMAX™ Supplement medium containing 10% FBS in each well of a 96-well plate for designated time points (24h, 48h, and 72h). The plates were then incubated at 37°C in a humidified CO_2_ incubator. At each time point, dye solution was added to each well, and the cells were incubated at 37°C for 4 hours, shielded from light. Following this incubation, stop solution was added, and the cells were further incubated for 1 hour at 37°C. Absorbance was subsequently measured at 570 nm using the AGILENT Biotek Synergy H1 multiplate reader.

### Migration assay

To assess cell migration, 10,000 cells were seeded into a 12-well insert (cellQART®, #9318002) with an 8µm pore size containing 350µl of FBS-free medium. Chemoattractant-containing medium, consisting of DMEM/F-12 with GlutaMAX™ Supplement and 10% FBS, was added to the bottom well. The cells were incubated for 48 hours at 37°C in a humidified CO_2_ incubator. After incubation, the medium was removed from the insert and non-migrated cells were gently swabbed from the top and bottom of each insert. The inserts were then fixed with 4% PFA for 15 minutes and stained with Crystal Violet (Sigma-Aldrich, #C0775) for 10 minutes. Once dried, the stain was solubilized in acetic acid solution (Supelco®, #5330010050) and the absorbance was read at 590 nm using the AGILENT Biotek Synergy H1 multiplate reader.

### RNA extraction, sequencing, and analysis

Total RNA was extracted using the NZY miRNA Isolation & RNA Clean-up Kit (NZYtech, #MB13402), following the manufacturer’s protocol. The purity and concentration of RNA from each sample were assessed using an Agilent TapeStation with an RNA ScreenTape Analysis Kit. mRNA enrichment was subsequently performed using polyA-tail-connected magnetic beads and Oligo. This was followed by double-stranded DNA synthesis and PCR amplification with specific primers. The PCR products were then subjected to thermal denaturation to produce single-stranded DNA, which was cyclized into a circular DNA library using bridge primers. Sequencing was conducted on the DNBSEQ platform (BGI Genomics Co.). Sequencing raw data were preprocessed using SOAPnuke software (BGI Genomics Co.). This involved removing reads with adaptor contamination, more than 5% N content, and low-quality reads (where more than 20% of bases had a quality score below 15). The resulting “Clean Reads” were saved in FASTQ format. RNA-seq data were analyzed using the Galaxy workbench platform [34], adhering to specific recommendations [35]. Quality controls and trimming of the reads were performed using MultiQC [36] and Cutadpt [37]. The reads were then mapped to the reference genome (Hg38, human genome build 38) using STAR [38]. From the mapped sequences, the number of reads per annotated gene was counted using featureCounts [39]. Subsequently, DESeq2 [40] was used to normalize the read counts and identify differentially expressed genes. Functional enrichment analysis of the differential expressed genes was performed using GSEA [41].

### *In vitro* drug response assay

5,000 cells were seeded in 96-well plates and incubated at 37°C in a humidified CO_2_ incubator. After 24 hours, the cells were treated with various concentrations of dasatinib (Selleckchem, #S1021) and incubated for an additional 72 hours at 37°C. The cells were then fixed with 0,2% glutaraldehyde for 10 minutes and stained with crystal violet for one hour at room temperature with shaking. Following staining, the plates were dried, and the dye was eluted with acetic acid. The absorbance was measured at 590 nm with the AGILENT Biotek Synergy H1 multiplate reader.

### Tumor growth and drug response in vivo

Female immune-compromised NMRI-Foxn1^nu/nu^ mice (Janvier Labs), typically 6 weeks old, were used to generate xenografts. Fresh PDX-derived tumor fragments or Hs578T cells (CONTROL and *TAGLN*-KO) were implanted subcutaneously or orthotopically, respectively. Mice were then treated daily with 20 mg/kg of dasatinib or vehicle (30% PEG and 5% Tween-80 in physiological serum) intraperitoneally. Treatment groups were randomized, and the health of the mice was monitored regularly. Tumor sizes were measured daily using digital calipers once growth was observed.

## Acknowledgements

VR is funded by RYC2018-024099-I. DOA is funded by PFERO2021.01. This work was funded by LABAE211626RODI and PID2020-114647RA-I00. We thank Comparative Medicine and Bioimage Centre of Catalunya (CMCiB), Confocal microscopy and Immunohistochemistry Unit (IIB Sant Pau), Microscopy Unit (IJC), and BIOBANK IGTP-HUGTP for their technical support.

