## Supplemental Figures and Tables for "Discovery and evaluation of novel biomarkers reveal dasatinib as a potential treatment for a specific subtype of Triple-Negative Breast Cancer"

### Supplemental Figure 1

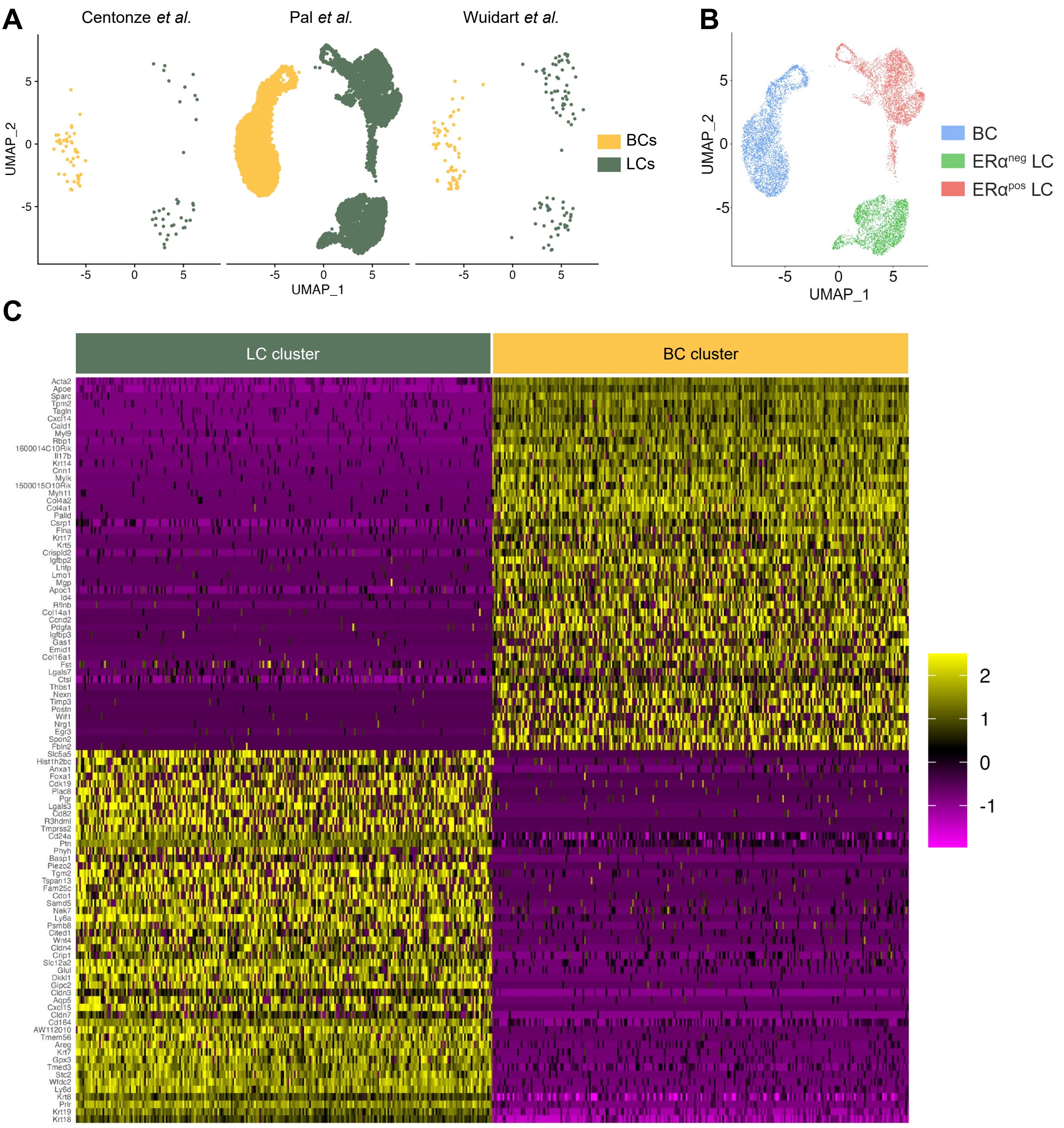


**Supplemental Figure 1 | Integration of three scRNA-seq datasets from adult murine mammary gland. A,** UMAP plots illustrating cell population distribution from three independent studies: Centonze et al. [21], Pal et al. [20], and Wuidart et al. [19]. Each point represents a single cell, color-coded to identify two cell compartments: BCs (dark green) and LCs (yellow). **B**, Integrated UMAP plot showing cell clusters from the three indicated scRNA-seq datasets. Color coding differentiates between three mammary epithelial cell types: BCs (blue), ERα-positive LC (red) and ERα-negative LCs (green). **C**, Heatmap depicting the differential gene expression profile of the Top 50 LC-associated and BC-associated genes. Each row corresponds to a gene, and each column to a single cell. Color intensity indicates normalized gene expression levels, ranging from -1 (purple) to +2 (yellow).

### Supplemental Figure 2

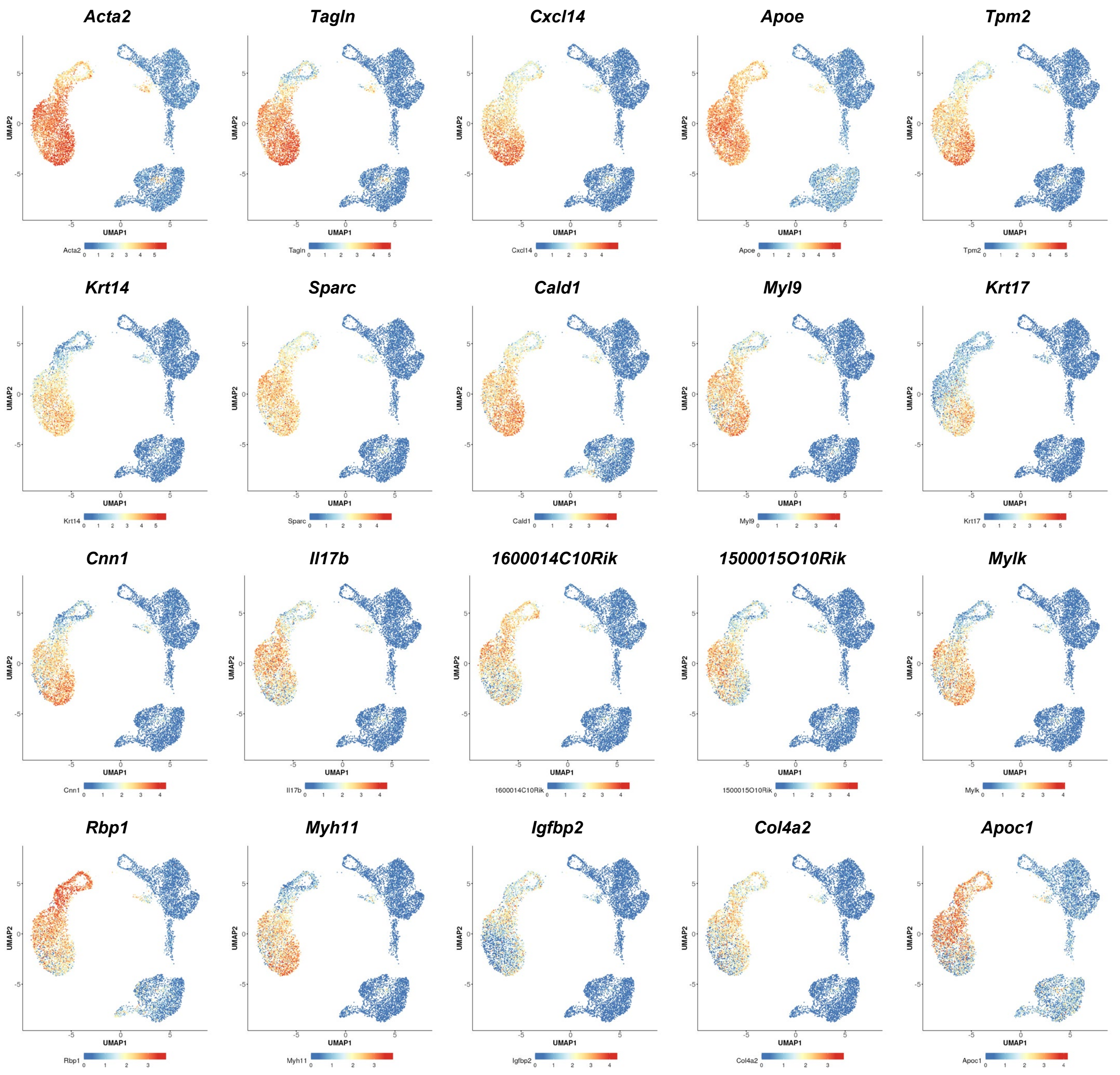


**Supplemental Figure 2 | Expression of BC-associated genes in murine mammary epithelial subpopulations.** This figure presents a series of UMAP plots illustrating the expression patterns of the top 20 BC-associated markers across integrated scRNA-seq data sets. Each plot corresponds to a specific gene, as indicated at the top of each plot. Points within the plots represent individual cells, with the color gradient indicating gene expression levels from low (blue) to high (red).

### Supplemental Figure 3

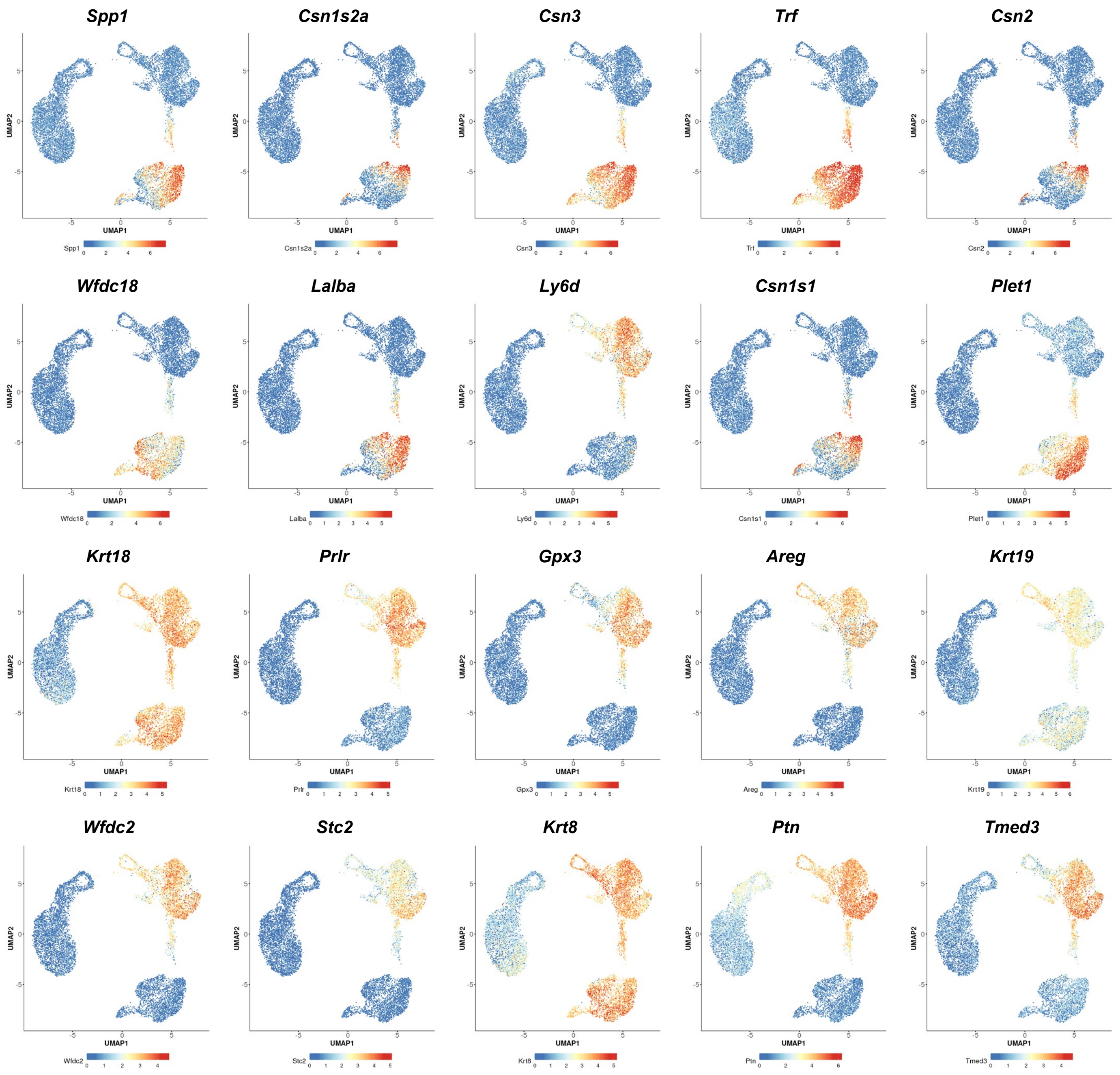


**Supplemental Figure 3 | Expression of LC-associated genes in murine mammary epithelial subpopulations.** This figure displays a series of UMAP plots depicting the expression patterns of the top 20 LC-associated markers across integrated scRNA-seq datasets. Each plot corresponds to a specific gene, as labeled at the top of each plot. Points within the plots represent individual cells, with the color gradient indicating gene expression levels ranging from low (blue) to high (red).

### Supplemental Figure 4

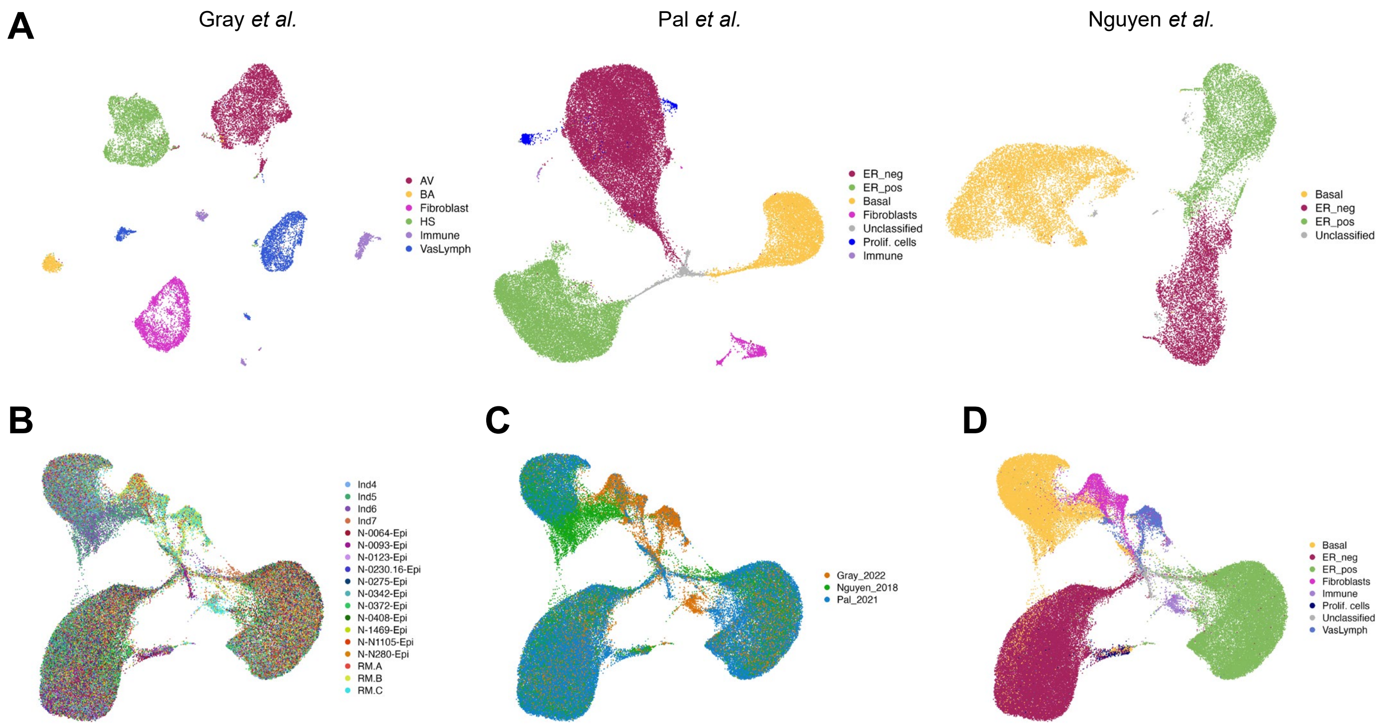


**Supplemental Figure 4 | Integration of three scRNA-seq datasets from healthy human mammary gland. A,** UMAP plots showing cell population distributions from three independent studies, labeled as Pal et al. [16], Gray et al. [23], and Nguyen et al. [15]. Each plot displays unique cell clusters with distinct expression profiles, categorized into cell types including immune cells, fibroblasts, and epithelial cells. **B-D**, Integrated UMAP plots combining data from all studies, with each plot distinctly color-coded according to the individual from whom the sample was taken (**B**), the origin of the study (**C**), and cell type classification (**D**).

### Supplemental Figure 5

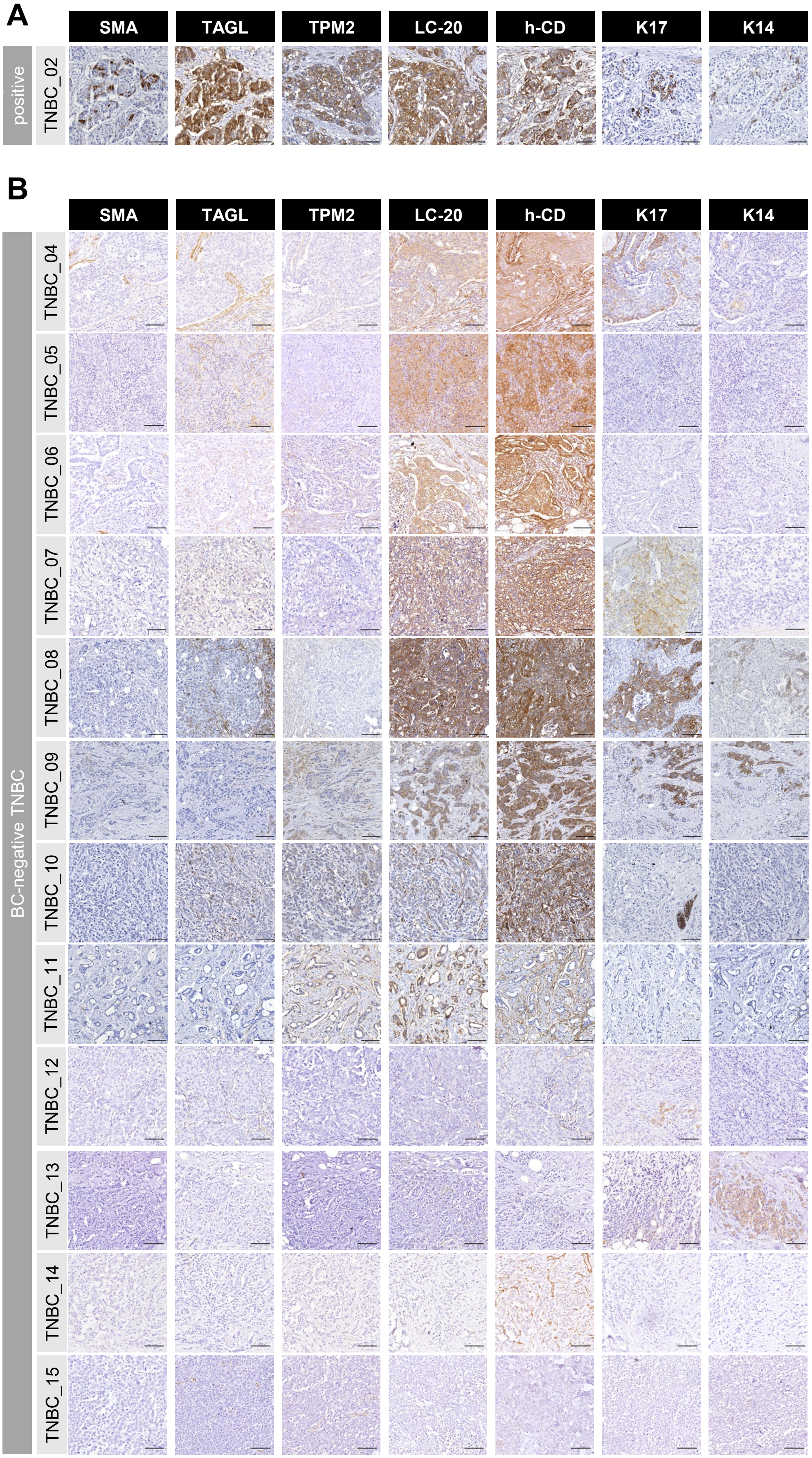


**Supplemental Figure 5 | Immunohistochemical validation of BC-markers in TNBC samples.** This figure presents representative sections of TNBC samples, comparing BC-positive (**A**) and BC-negative (**B**) categories. Each row corresponds to a specific patient sample, labeled from TNBC-02 to TNBC-15. Columns represent staining for various BC-markers: SMA, TAGL, TPM2, LC-20, h-CD, K17, and K14. Staining intensity varies, with the color intensity indicative of marker expression. Nuclei are counterstained with hematoxylin. Scale bar represents 100 µm.

### Supplemental Figure 6

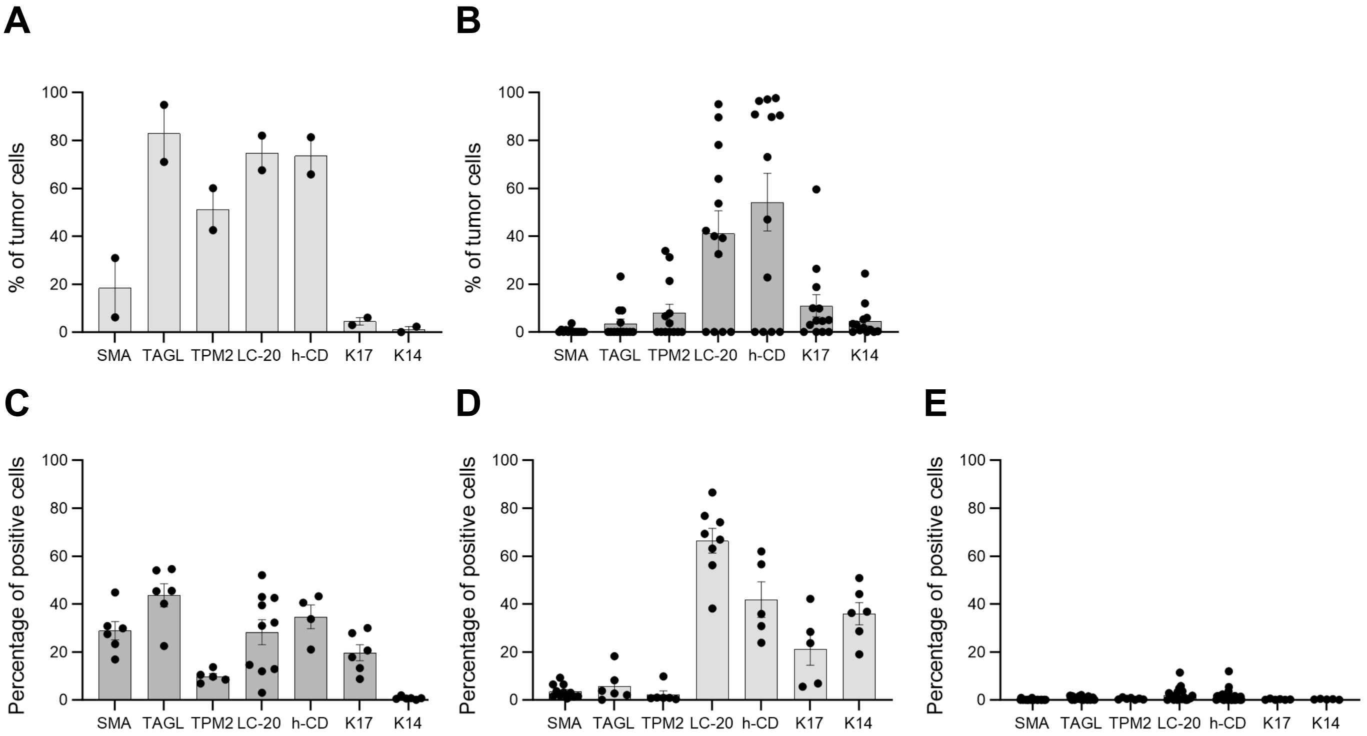


**Supplemental Figure 6 | Immunohistochemistry quantifications of BC-markers in TNBC and HR-positive samples.** This figure shows bar plots of the percentage of tumor cells expressing each BC-marker across various sample types: **A**, percentage of tumor cells positive for each BC-marker in BC-positive TNBC samples. **B**, percentage of tumor cells positive for each BC-marker in BC-negative TNBC samples. **C,** Percentage of tumor cells positive for each BC-marker per tumor area in low-grade DCIS. **D**, percentage of tumor cells positive for each BC-marker per tumor area in DCIS. **E**, percentage of tumor cells positive for each BC-marker per tumor area in IDC. Each bar represents the mean percentage of positive cells, with individual data points shown. Error bars represent standard error of the mean.

### Supplemental Figure 7

**A**

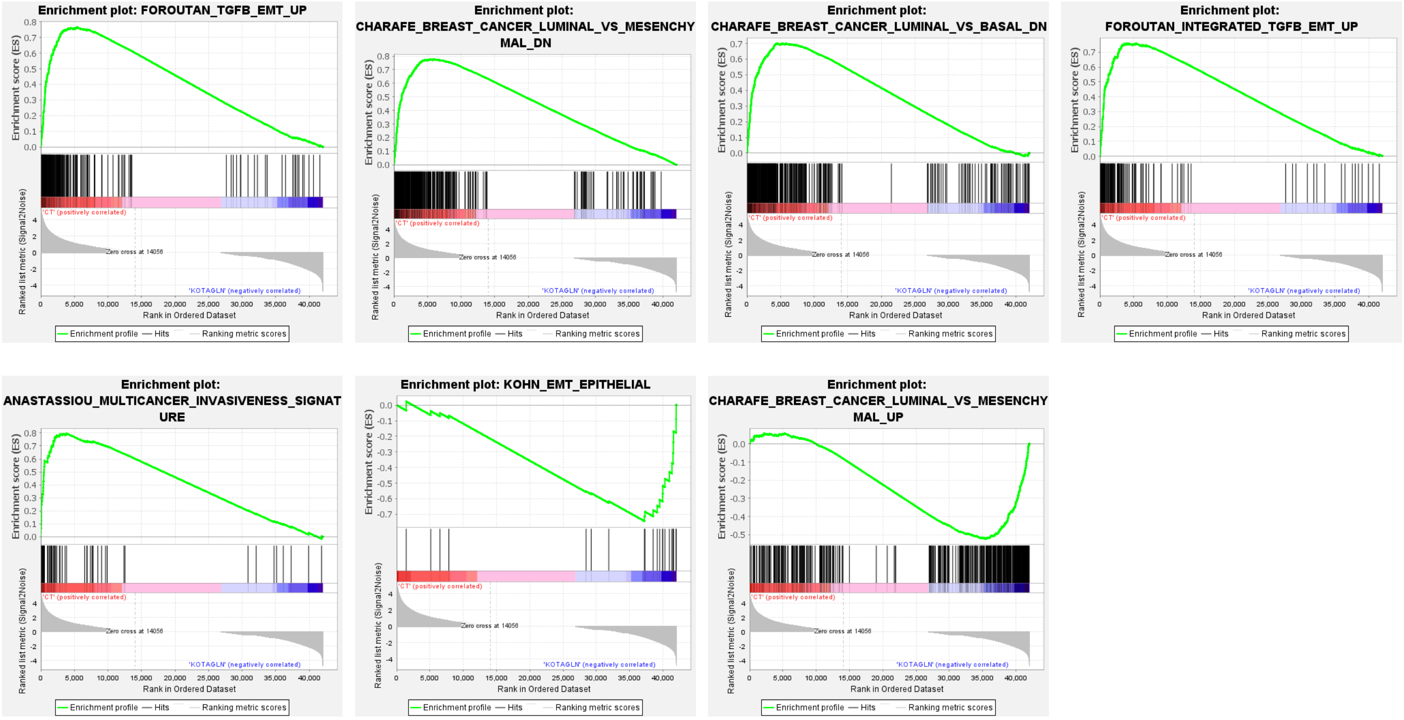


**B**

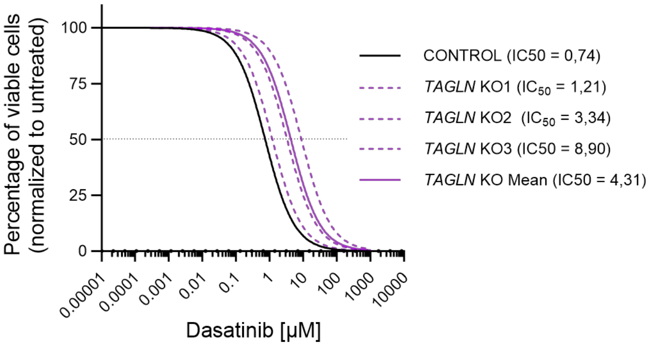


**Supplemental Figure 7 | Functional validation of TAGLN-KO cells. A,** Enrichment plots of the GSEA for epithelial to mesenchymal transition in CONTROL versus TAGLN-KO cells. **B**, Drug-response curves for cell viability of BT-549 cells, comparing CONTROL (black) and TAGLN-KO (purple) treated with increasing concentrations of dasatinib. Solid lines represent the mean of three biological replicates, performed in technical replicates, and error bars indicate ±SEM. Dashed lines indicate the $IC_{50}$ for each condition.

### Supplemental Figure 8

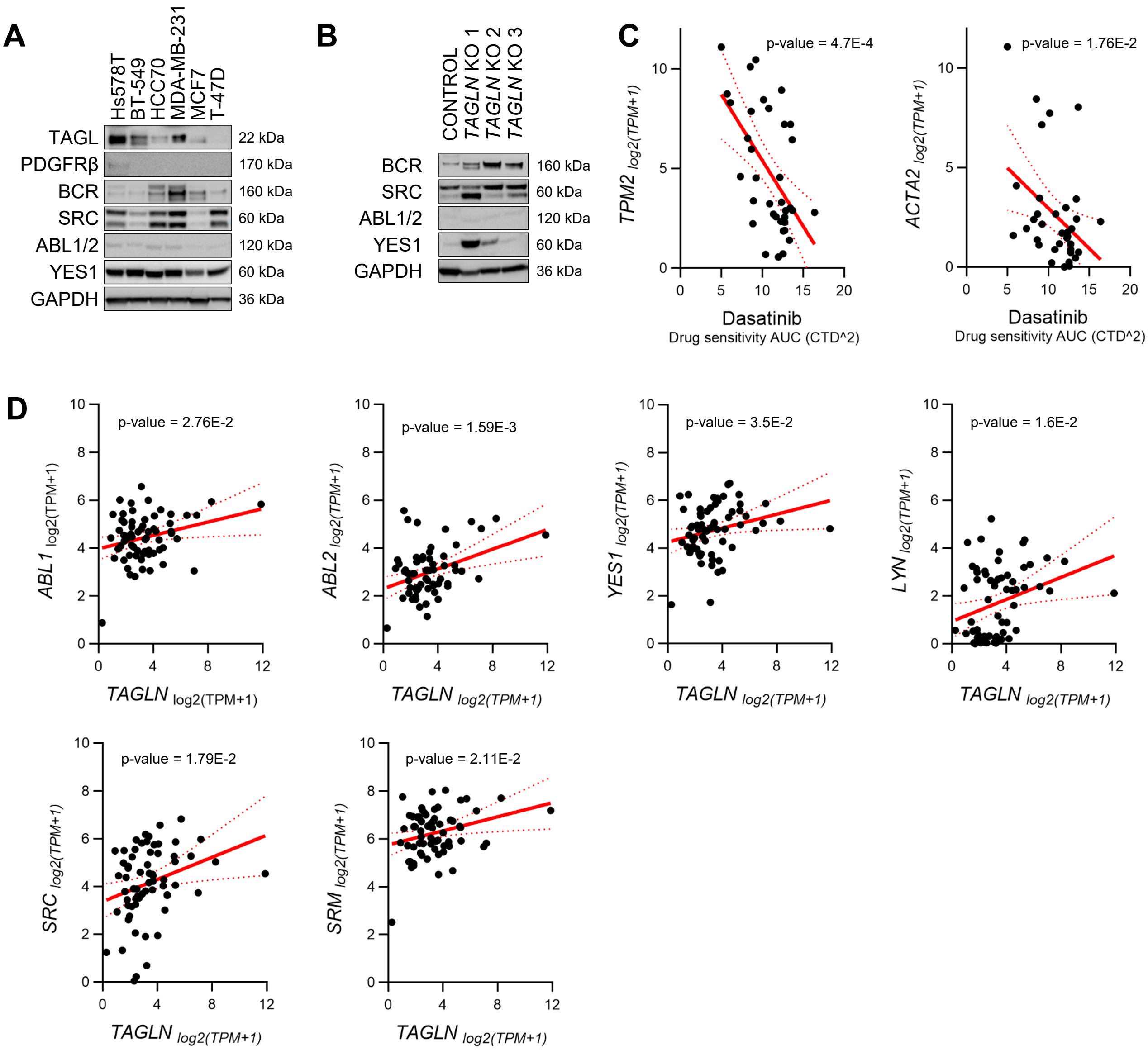


**Supplemental Figure 8 | Analysis of *TAGLN* expression and dasatinib target interaction in cancer cell lines**. **A,** WB analysis showing protein levels of TAGL and indicated dasatinib targets in breast cancer cell lines. **B,** WB analysis showing protein levels of indicated dasatinib targets in Hs578T CONTROL and *TAGLN*-KO cells. **C,** Scatter plots illustrating the correlation between dasatinib sensitivity (measured as AUC) and the expression of *TPM2* and *ACTA2*, across various breast cancer cell lines. Each dot represents a cell line with dasatinib sensitivity on the X-axis and gene expression levels on the Y-axis. Red dashed lines indicate the regression lines, and the associated p-values are displayed to show statistical significance. **D,** Scatter plots displaying the correlation between *TAGLN* expression and the expression of dasatinib target genes (*ABL1, ABL2, YES1, LYN, SRC*, and *SRM*) in breast cancer cell lines. Each plot shows the expression of *TAGLN* (log2(TPM+1)) on the X-axis against the expression of a target gene (log2(TPM+1)) on the Y-axis. Data points are depicted as black dots, with red dashed lines indicating the trend lines and p-values provided for each correlation.

### Supplemental Table 1

| Target | Reference | Supplier | [IHC] | [IF] | [WB] |
|---|---|---|---|---|---|
| Smooth Muscle Actin Anticuerpo (B4) | sc-53142 | Santa cruz | 2 ug/ml | | |
| Transgelin Anticuerpo (6G6) | sc-53932 | Santa cruz | 2 ug/ml | 2 ug/ml | 0,2 ug/ml |
| CXCL14 Polyclonal Antibody | PA5-106402 | Invitrogen | 2 ug/ml | | |
| Tropomyosin Anticuerpo (F-6) | sc-74480 | Santa cruz | 1 ug/ml | | |
| Anti-KRT14 antibody | HPA023040 | Sigma-Aldrich | 0,1 mg/ml | | |
| SPARC Anticuerpo (AON-1) | sc-33645 | Santa cruz | 2 ug/ml | | |
| Anti-CALD1 antibody | HPA017330 | Sigma-Aldrich | 0,7 mg/ml | | |
| MYL9/MYL12A/B Anticuerpo (E-4) | sc-28329 | Santa cruz | 1 ug/ml | | |
| Cytokeratin 17 Anticuerpo (E-4) | sc-393002 | Santa cruz | 2 ug/ml | | |
| Apolipoprotein E/apoE Anticuerpo (WU E-4) | sc-53570 | Santa cruz | 2 ug/ml | | |
| BCR Polyclonal Antibody | 11592600 | Invitrogen | | | 0,45 ug/ml |
| Rabbit anti-PDGFR beta Recombinant Monoclonal Antibody [BLR081G] | A700-081-T | Bethyl Laboratories | | | 1 ug/ml |
| SRC Monoclonal Antibody (184Q20) | 10776283 | Invitrogen | | | 0,5 ug/ml |
| GAPDH Anticuerpo (0411) | sc-47724 | Santa cruz | | | 0,2 ug/ml |
| YES1-Specific Polyclonal antibody | 20243-1-AP-150UL | LABCLINICS SA | | | 1,6 ug/ml |
| ABL1/ABL2 Poyconcal antibody | PA5-114805 | INVITROGEN | | | 1 ug/ml |

**Table S1 | Primary antibodies used for immunohistochemistry (IHC), immunofluorescence (IF), and Western blot (WB) assays.** This table lists the antibodies utilized across various assays in the study. Details include the target protein, the catalog number, supplier, and the specific concentrations used for each assay type (IHC, IF, WB). Concentrations are provided in micrograms per milliliter (µg/ml) or milligrams per milliliter (mg/ml) as appropriate. Antibodies are grouped by their respective applications, providing a comprehensive resource for replication of the experimental conditions.

### Supplemental Table 2

| Target | Reference | Supplier | [IHC] | [IF] | [WB] |
|---|---|---|---|---|---|
| Goat anti-Mouse IgG (H+L) Secondary Antibody, HRP | 31430 | Thermo Scientific | 3,2 ug/ml | | 0,1 ug/ml |
| Goat anti-Rabbit IgG (H+L) Secondary Antibody, HRP | 31460 | Thermo Scientific | 3,2 ug/ml | | 0,1 ug/ml |
| Alexa Fluor® 488 AffiniPure Donkey Anti-Mouse IgG (H+L) | 715-545-150 | Jackson Immunoresearch | | 1 ug/ml | |

**Table S2 | Secondary antibodies used for immunohistochemistry (IHC), immunofluorescence (IF), and Western blot (WB) assays.** This table lists the antibodies utilized across various assays in the study. Details include the target protein, the catalog number, supplier, and the specific concentrations used for each assay type (IHC, IF, WB). Concentrations are provided in micrograms per milliliter (µg/ml) or milligrams per milliliter (mg/ml) as appropriate. Antibodies are grouped by their respective applications, providing a comprehensive resource for replication of the experimental conditions.
